# Promotech: A general tool for bacterial promoter recognition

**DOI:** 10.1101/2021.07.16.452684

**Authors:** Ruben Chevez-Guardado, Lourdes Peña-Castillo

## Abstract

Promoters are genomic regions where the transcription machinery binds to initiate the transcription of specific genes. Computational tools for identifying bacterial promoters have been around for decades. However, most of these tools were designed to recognize promoters in one or few bacterial species. Here, we present Promotech, a machine-learning-based method for promoter recognition in a wide range of bacterial species. We compared Promotech’s performance with the performance of five other promoter prediction methods. Promotech outperformed these other programs in terms of area under the precision-recall curve (AUPRC) or precision at the same level of recall. Promotech is available at https://github.com/BioinformaticsLabAtMUN/PromoTech.

## Background

Promoters are DNA segments essential for the initiation of transcription at a defined location in the genome, which are recognized by a specific RNA polymerase (RNAP) holoenzyme (E*σ*) [1]. E*σ* is formed by RNAP and a *σ* factor. *σ* factors are bacterial DNA binding regulatory proteins of transcription initiation that enable specific binding of RNAP to promoters [1]. Recognizing promoters is critical for understanding bacterial gene expression regulation. There have been numerous bioinformatics tools developed to recognize bacterial promoter sequences (Supplementary Table 1). However, most of these tools were designed to recognize promoters in *Escherichia coli* or in few (2 or 3) bacterial species, and their applicability to a wider range of bacterial species is unproven. Additionally, the performance of current tools rapidly decreases when applied to whole genomes and thus it is common practice to restrict the size of the input sequence to a few hundred nucleotides.

Shahmuradov et al. [2] evaluated the performance of their method (bTSSfinder) and other three methods on ten bacterial species belonging to five different phyla. The best average sensitivity (recall) values obtained were 59% and 49% by bTSSfinder and BPROM [3], respectively; while, bTSSfinder achieved higher accuracy than the other three assessed tools. These results are promising as they showed that it is possible to recognize promoters of several bacterial species even when the methods were designed for specific bacterial species. BPROM uses five relatively conserved motifs from *E. coli* to identify promoters, and bTSSfinder focuses on *E. coli* and three species of Cyanobacteria. Based on this, we hypothesized that predictive performance can be improved if a method is trained on data from a diverse group of bacterial species.

Another bacterial promoter detection method evaluated on a multi-species data set is G4PromFinder [4]. G4PromFinder utilizes conserved motifs and focuses on *Streptomyces coelicolor* A3(2) and *Pseudomonas aeruginosa* PA14. Di Salvo et al. comparatively assessed G4PromFinder’s performance in terms of F1-score with that of bTSSfinder, PePPER [5] and PromPredict [6, 7], and found that G4PromFinder outperformed the other three tools in GC rich bacterial genomes. There are several recently published *E. coli* promoter prediction methods such as MULTiPly [8], SELECTOR [9], iPromoter-BnCNN [10], IBPP [11], iPromoter-2L [12], among others (Supplementary Table 1). Cassiano and Silva-Rocha [13] carried out a comparative assessment of bacterial promoter prediction tools for identifying *E. coli σ*^70^ promoters. In their benchmark, their found that iPro70-FMWin [14] achieved the best results in terms of accuracy and MCC.

Here, we developed a general (species independent) bacterial promoter recognition method, Promotech, trained on a large data set of promoter sequences of nine distinct bacterial species belonging to five different phyla (namely, Actinobacteria, Chlamydiae, Firmicutes, Proteobacteria, and Spirochaetes). As promoters are typically located directly upstream of the transcription start site (TSS), we used published TSS global maps obtained using sequencing technology such as dRNA-seq [15] and Cappable-seq [16] to define promoter sequences. We trained and evaluated twelve Random Forests and Recurrent Neural Networks models using these data to select Promotech’s classification model (Fig. 1). Finally, we compared the performance of Promotech with that of five other bacterial promoter prediction methods on independent data from four bacterial species. Promotech outperformed all these five methods in terms of the area under the ROC curve (AUROC), the area under the precision-recall curve (AUPRC) and precision at a specific recall level.

**Figure 1.**
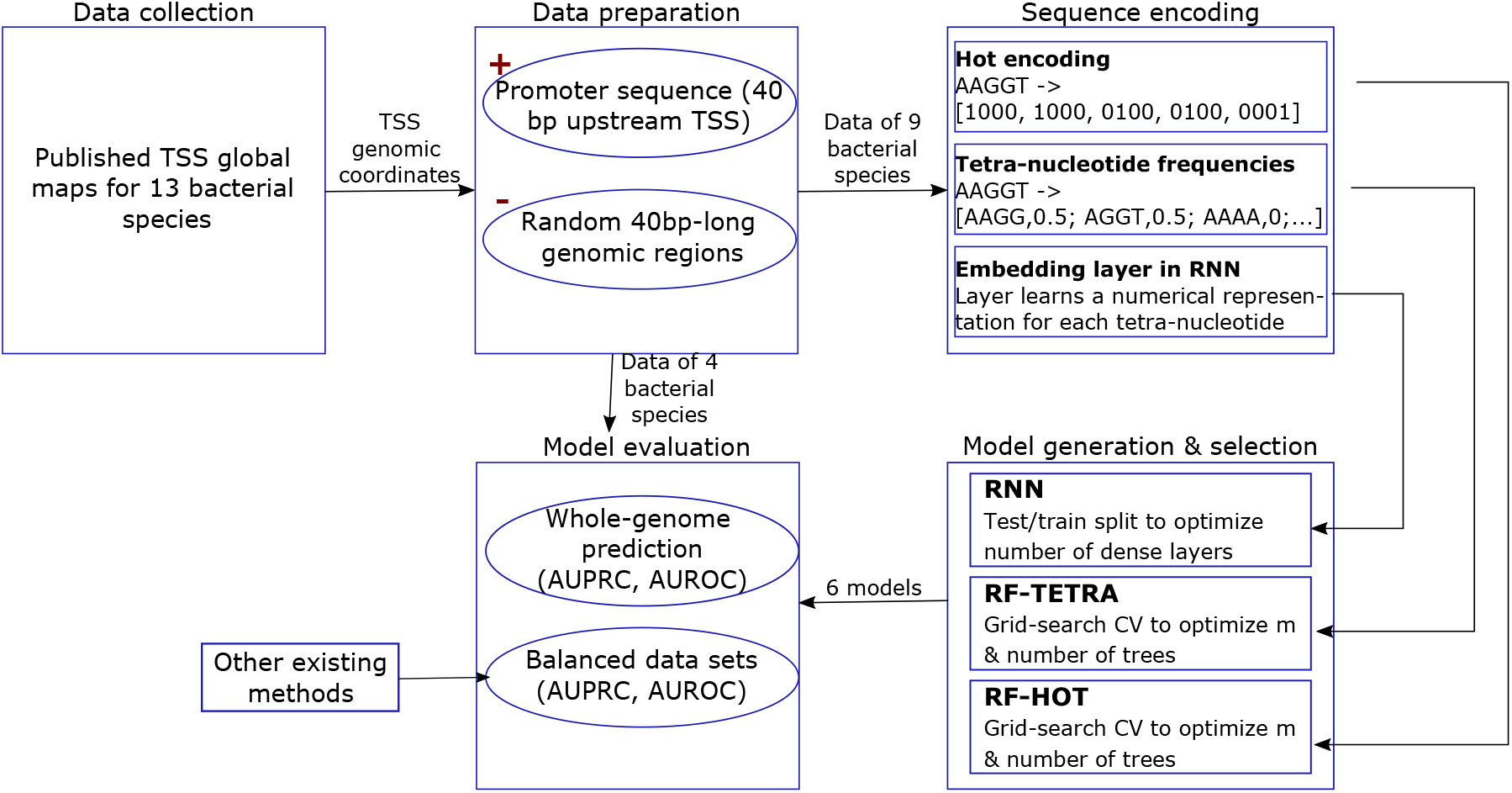
Flowchart illustrating our methodology.

## Results and Discussion

### Variety of training and validation data

We obtained a large amount of promoter sequences from published global TSS maps (Supplementary Tables 2). On both the training and the validation data, we had bacterial species belonging to distinct phyla and having a wide range (from 30% to 72%) of GC content (Tables 1 and 2). In total, our training data contained 27,766 promoter sequences, and our validation data contained 11,615 promoter sequences (Supplementary Tables 3 and 4).

**Table 1.** Training dataset’s characteristics.

| Bacterial species | Phylum | GC content (%) |
| --- | --- | --- |
| *Streptomyces coelicolor* | Actinobacteria | 71.98 |
| *Chlamydia pneumoniae* | Chlamydiae | 40.6 |
| *Streptococcus pyogenes* | Firmicutes | 38.4 |
| *Salmonella enterica*<br>serovar Typhimurium | Proteobacteria | 52.1 |
| *Escherichia coli* | Proteobacteria | 50.6 |
| *Shewanella oneidensis* | Proteobacteria | 46 |
| *Helicobacter pylori* | Proteobacteria | 38.9 |
| *Campylobacter jejuni* | Proteobacteria | 30.4 |
| *Leptospira interrogans* | Spirochaetes | 35 |

**Table 2.** Validation dataset’s characteristics.

| Bacterial species | Phylum | GC content (%) |
| --- | --- | --- |
| *Mycobacterium smegmatis* | Actinobacteria | 67.4 |
| *Bacillus amyloliquefaciens* | Firmicutes | 46.4 |
| *Lachnoclostridium*<br>*phytofermentans* | Firmicutes | 35.6 |
| *Rhodobacter capsulatus* | Proteobacteria | 66.5 |

**Table 3.** The AUPRC and AUROC obtained in 25% of the training dataset left out for testing. The dataset used has a 1:10 ratio of positive to negative instances. The numbers in bold indicate the models with the highest AUPRC / AUROC per machine learning method.

| MODELS | AUPRC | AUROC |
| --- | --- | --- |
| RF-HOT | **0.802** | **0.938** |
| RF-TETRA | 0.593 | 0.844 |
| GRU-0 | 0.752 | 0.929 |
| GRU-1 | **0.778** | **0.934** |
| GRU-2 | 0.753 | 0.929 |
| GRU-3 | 0.728 | 0.922 |
| GRU-4 | 0.728 | 0.923 |
| LSTM-0 | 0.734 | 0.923 |
| LSTM-1 | 0.744 | 0.927 |
| LSTM-2 | 0.739 | 0.923 |
| LSTM-3 | **0.748** | 0.924 |
| LSTM-4 | **0.748** | **0.928** |

**Table 4.** Impurity-based feature importance ranking generated by the RF-TETRA model.

| Ranking | Tetra-nucleotide | Score |
| --- | --- | --- |
| 1 | TATA | 0.023±0.015 |
| 2 | ATAA | 0.014±0.009 |
| 3 | TAAT | 0.014±0.008 |
| 4 | TTAT | 0.011±0.007 |
| 5 | AAAA | 0.010±0.001 |
| 6 | TTTT | 0.010±0.001 |
| 7 | GTTA | 0.009±0.004 |
| 8 | TATT | 0.009±0.004 |
| 9 | TAAA | 0.009±0.002 |
| 10 | AATA | 0.008±0.004 |

### Model selection

To select Promotech’s classification model, we built two and ten models using Random Forest (RF) [17, 18] and Recurrent Neural Networks (RNN) [19], respectively (Fig. 1). The RF models consisted of one trained with hot-encoded features (RF-HOT) and another trained with tetra-nucleotide frequencies (RF-TETRA). To calculate the tetra-nucleotide frequencies vector of a given sequence, the number of occurrences of each possible 4-nucleotide DNA sequence (4-mer) in that sequence is divided by the total number of 4-mers in it. The optimal parameters of the RF models were selected using 10-fold grid-search cross-validation on 75% of the training data. The RNN models consisted of five Long Short Term Memory (LSTM) [20] and five Gated Recurrent Unit (GRU) [21] models having zero to four hidden layers, and a word embedding [22] layer to obtain a numerical representation of the promoter sequences (Table 5 in Additional File 1). From now on these RNN models are denoted as GRU-X or LSTM-X, where X indicates the number of hidden layers. All the models were trained using an unbalanced dataset with a 1:10 ratio of positive to negative instances to simulate the small number of promoters in a whole bacterial genome. Due to time constraints we were unable to run grid-search cross-validation for the RNNs, and assessed these models by randomly splitting the training data into 75% for training and 25% for testing. The best performing models per machine learning method in terms of AUPRC and AUROC were RF-HOT, GRU-1 and LSTM-4, as shown in Table 3. RF-HOT was the model with the highest AUPRC overall.

**Table 5.** Permutation-based feature importance ranking generated by the RF-TETRA model.

| Ranking | Tetra-nucleotide | Score |
| --- | --- | --- |
| 1 | ATAA | 0.052±0.001 |
| 2 | TATA | 0.048±0.001 |
| 3 | TAAT | 0.046±0.002 |
| 4 | TTAT | 0.039±0.001 |
| 5 | GTTA | 0.036±0.001 |
| 6 | TAAA | 0.035±0.001 |
| 7 | AATA | 0.035±0.001 |
| 8 | ATTA | 0.033±0.001 |
| 9 | TATT | 0.031±0.001 |
| 10 | AATT | 0.031±0.000 |

### Model interpretation

To interpret the models created, we performed feature importance analysis to find out motifs recognized by the models. To do this, we obtained the feature importance ranking from the RF models. First the importance scores were calculated using permutation-based importance (also called mean decrease in accuracy) [23] and impurity-based importance [18] on RF-TETRA. The most important tetramers based on the impurity-based importance score from RF-TETRA were TATA, ATAA, TAAT, TTAT, AAAA, and TTTT. The test was repeated using the permutation-based importance score and permuting each feature five times. Both tests produced similar results having the same tetra-nucleotide sequences appearing at the top of the ranking, only varying their relative ranking position and score (Tables 4 and 5).

The feature importance analysis was repeated on the RF-HOT model. In RF-HOT, each feature represents the presence of one of the four possible nucleotides: adenine (A), thymine (T), guanine (G), and cytosine (C) for the current position in the range of -39 to 0 relative to the TSS. Each nucleotide was represented as a 4-digit binary number, i.e., A (1000), G (0100), C (0010), and T (0001). The permutation and impurity-based feature importance ranking generated by RF-HOT provided the most important positions in the range of -39 to 0 relative to the TSS and the nucleotide with the most relevance for each position. To have visual representations of these results, each nucleotide’s importance score was plotted on a bar graph (Figs. 2 and 3). These results suggest that having adenine (A) and thymine (T) in the range of -8 to -12 relative to the TSS is highly important for promoter recognition. Additionally these results suggest that the RF models learn to identify the Pribnow-Schaller box [24, 25], which is a six nucleotide consensus sequence (TATAAT), commonly located around ten base pairs upstream from the TSS.

**Figure 2.**
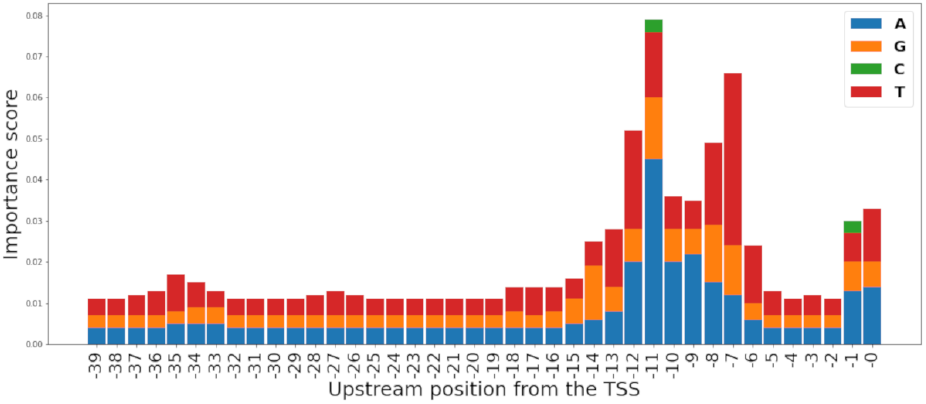
Impurity-based feature importance scores per nucleotide per position relative to the TSS as calculated from the RF-HOT model.

**Figure 3.**
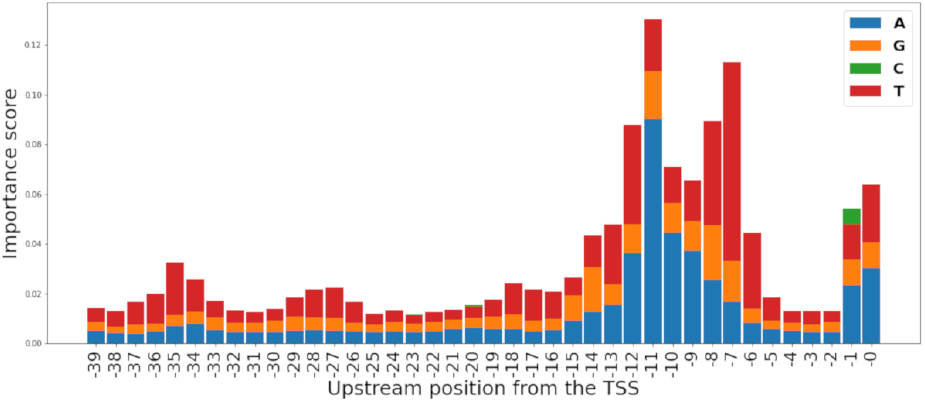
Permutation-based feature importance scores per nucleotide per position relative to the TSS as calculated from the RF-HOT model.

### Genome-wide promoter prediction assessment

We designed our first two assessments to demonstrate that Promotech was able to make predictions for a whole bacterial genome. Five models were selected for this assessment: the best models per machine learning method (RF-HOT, GRU-1, and LSTM-4), and GRU-0 and LSTM-3. GRU-0 and LSTM-3 were selected to evaluate the effect in performance if the model had one less hidden layer. As the models were trained on 40-nt long sequences, to do whole-genome predictions, we needed to cut the genome in 40-nt long sequences. Thus, we traversed each genome with a sliding window with a one nt step and a 40 nt window size. The sliced sequences were then pre-processed and fed to the model twice, first, using a forward strand configuration and then using a backwards strand configuration. These steps were repeated for each bacterium on the validation set; namely, *M. smegmatis, L. phytofermentans, B. amyloliquefaciens*, and *R. capsulatus*. This was a computationally demanding assessment; as, for example, the sliding window created 6,988,167 sequences of 40 nt when used on the *M. smegmatis* genome. Each sequence was given a second time with a backward strand configuration ending up with 13,976,336 sequences. Thus, each of the four selected models was executed roughly 14 million times for the *M. smegmatis* genome. The process took around 4 hours to run per bacterium per model, including the sliding window, data pre-processing, and model’s execution. Increasing the step size decreased the execution time but also decreased the model performance.

In the first assessment predicted promoters were considered true positives if they have at least 10% sequence overlap with an actual promoter. Table 6 shows the average AUPRC and AUROC obtained by each model. The PRC and ROC obtained per bacterium are shown in Supplementary Figs. 1-4 in Additional File 1.

**Table 6.** Average AUPRC and AUROC ± standard deviation obtained per model across the validation set when requiring that predicted promoters have at least 10% sequence overlap with the actual promoters to be considered true positives. The numbers in bold indicate the model with the highest performance.

| Model | Mean AUPRC | Mean AUROC |
| --- | --- | --- |
| RF-HOT | **0.143 ± 0.104** | **0.823 ± 0.160** |
| GRU-0 | 0.026 ± 0.016 | 0.687 ± 0.063 |
| GRU-1 | 0.025 ± 0.017 | 0.675 ± 0.048 |
| LSTM-3 | 0.034 ± 0.017 | 0.677 ± 0.019 |
| LSTM-4 | 0.033 ± 0.023 | 0.631 ± 0.040 |

RF-HOT achieved the best overall AUPRC (0.14) and AUROC (0.82) (Table 6). An AUPRC of 0.14 might seem low, but if one considers that there are millions of 40 nt-long sequences in a bacterial genome and only a few thousand of these sequences are actual promoters, then this performance is much better than random guessing. For example, *M. smegmatis* has four thousand actual promoters (Supplementary Table 2) and 14 million 40 nt-long genomic sequences, thus a random classifier has an average AUPRC of 0.0003. RF-HOT achieved an AUPRC of 0.27 in *M. smegmatis* genome which is roughly a thousand-fold improvement over random performance.

To gain insight into the behaviour of the models, we visually inspected the location of predicted promoters and observed that many predicted promoters were located nearby actual promoters (Fig. 4). To account for this, we re-evaluated the models performance to count as correct predictions those within 100nt of an actual promoter. We called this task “the cluster promoter prediction”. Assessing the performance of the models using the cluster promoter prediction method increased AUPRC 2 to 6 times and AUROC by 1.5 times the values obtained in the first assessment (Figs. 5, 6, 7, and 8, and Table 7). This suggests that our models predict promoters in the proximity of actual promoters but are unable to recognize the exact genomic location of the actual promoters.

**Table 7.** Average AUPRC and AUROC standard deviation obtained per model across the validation set on the cluster promoter prediction task. The numbers in bold indicate the highest AUPRC and AUROC.

| Model | Mean AUPRC | Mean AUROC |
| --- | --- | --- |
| RF-HOT | **0.708±0.182** | **0.939±0.050** |
| GRU-0 | 0.384±0.218 | 0.898±0.030 |
| GRU-1 | 0.365±0.219 | 0.894±0.027 |
| LSTM-3 | 0.404±0.199 | 0.900±0.011 |
| LSTM-4 | 0.379±0.200 | 0.867±0.033 |

**Figure 4.**
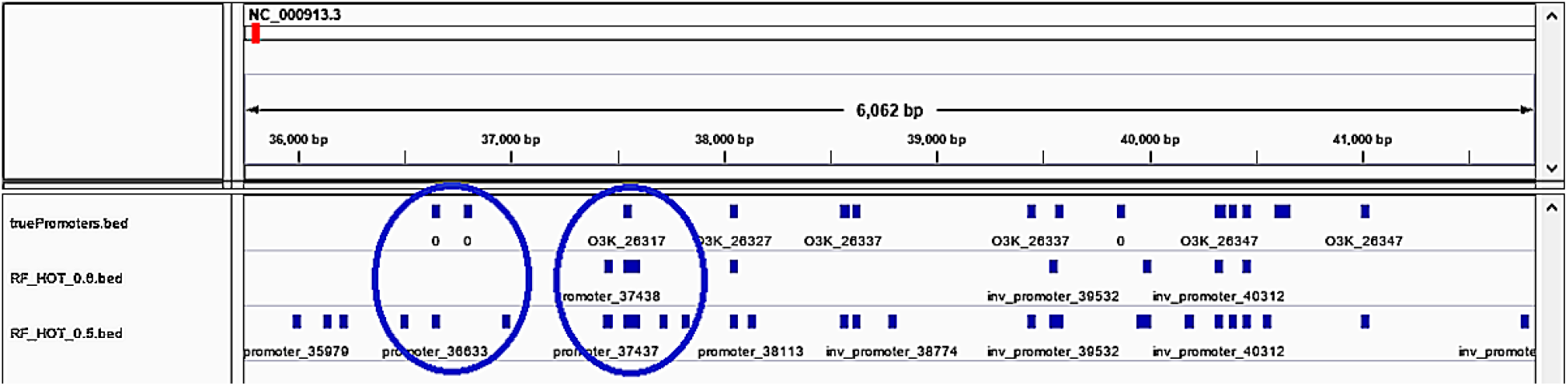
Predicted promoters observed in actual promoters’ proximity but not overlapping. Blue squares on the first row indicate the location of actual promoters while blue squares on the second and third rows indicate the location of predicted promoters with a predicted probability of 0.6 and 0.5 respectively. Within each circle a predicted promoter cluster is shown.

**Figure 5.**
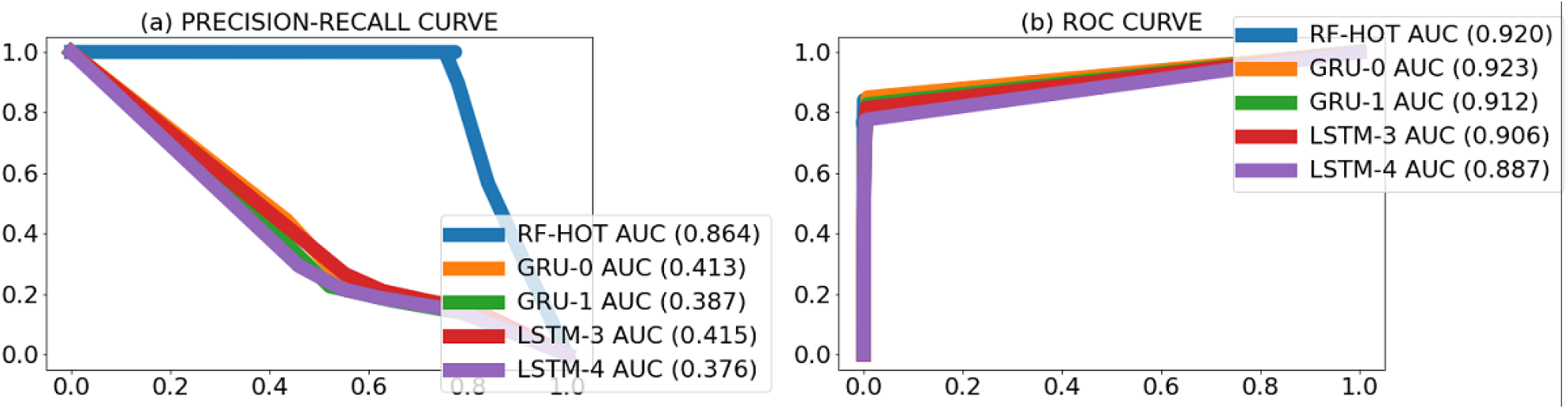
PR curves (a) and ROC curves (b) per model obtained when counting predicted promoters nearby actual promoters as true positives on *M. smegmatis* str. MC2 155 bacterium. Numbers between brackets beside the model ID indicate AUPRC (a) and AUROC (b) of that model.

**Figure 6.**
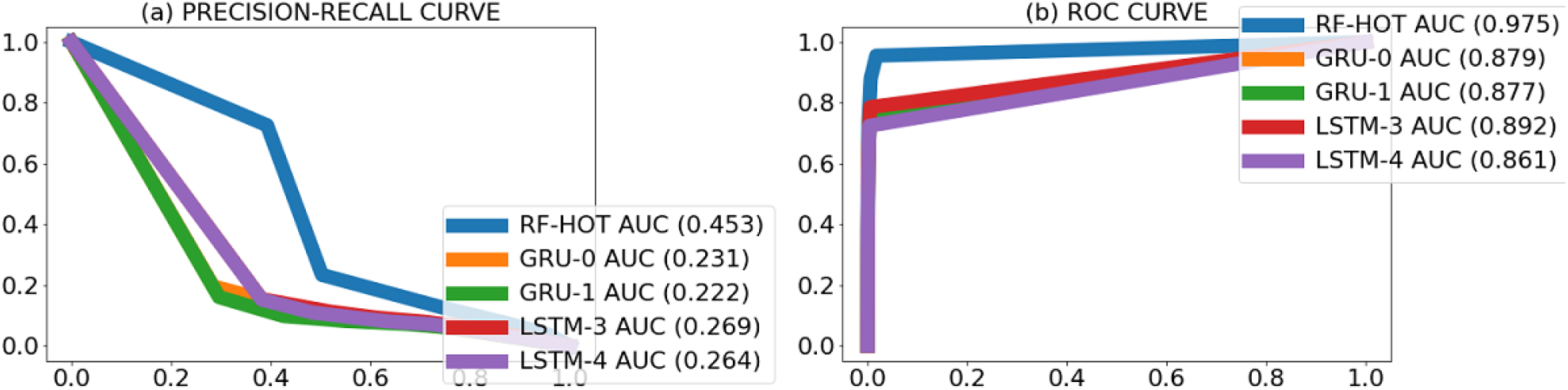
PR curves (a) and ROC curves (b) per model obtained when counting predicted promoters nearby actual promoters as true positives on *L. phytofermentans* ISDg bacterium. Numbers between brackets beside the model ID indicate AUPRC (a) and AUROC (b) of that model.

**Figure 7.**
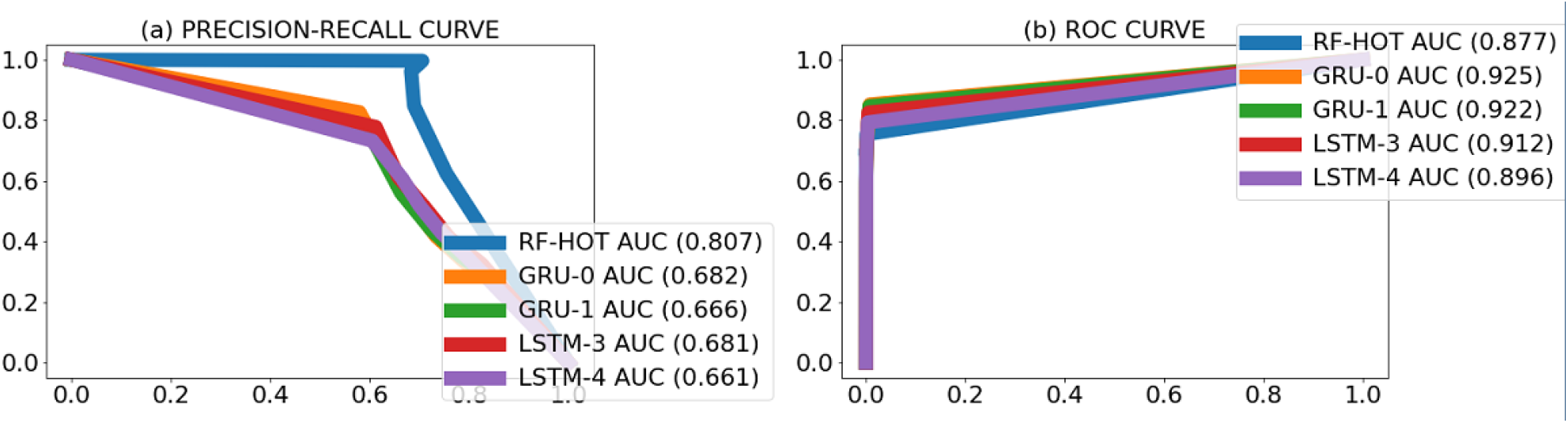
PR curves (a) and ROC curves (b) per model obtained when counting predicted promoters nearby actual promoters as true positives on *R. capsulatus* SB 1003 bacterium. Numbers between brackets beside the model ID indicate AUPRC (a) and AUROC (b) of that model.

**Figure 8.**
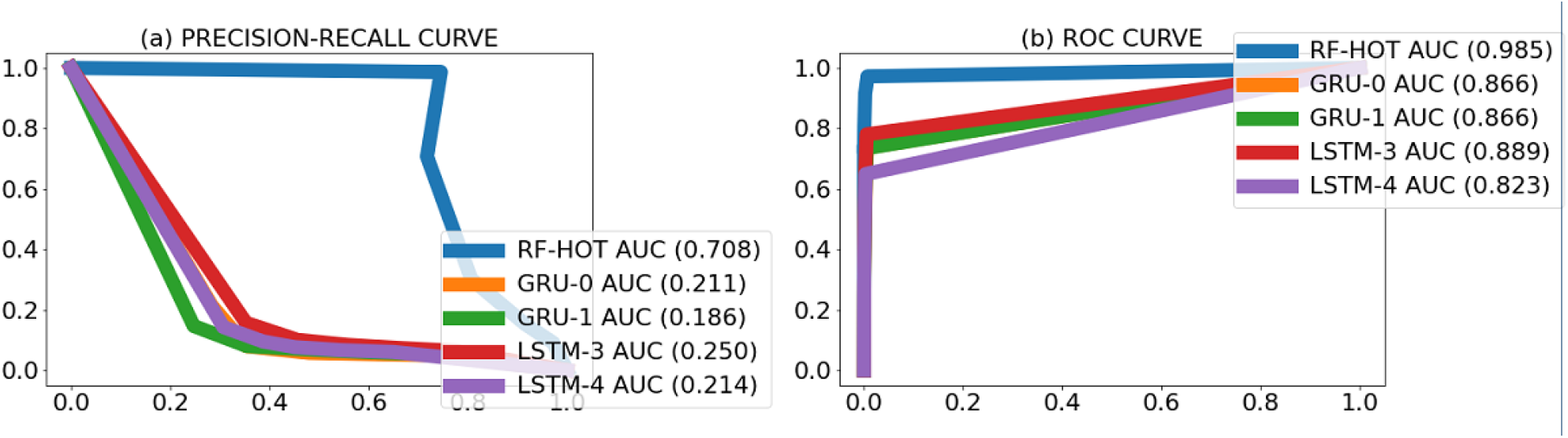
PR curves (a) and ROC curves (b) per model obtained when counting predicted promoters nearby actual promoters as true positives on *B. amyloliquefaciens* XH7 bacterium. Numbers between brackets beside the model ID indicate AUPRC (a) and AUROC (b) of that model.

### Performance comparison with existing methods

To compare Promotech’s performance with that of other existing methods, we used four independent test datasets containing promoters found by global TSS mapping using sequencing technologies. As in the previous assessments, the independent datasets included *B. amyloliquefaciens, L. phytofermentans, M. smegmatis* and *R. capsulatus*. On this assessment, the datasets have a 1:1 ratio of positive to negative instances (Supplementary Table 4). As these datasets contained thousands instead of millions of sequences, we were able to include the RF-TETRA model that failed to run on a whole genome (due to memory issues).

As Promotech’s goal is to be applicable to a wide range of bacterial species, we compared Promotech models with other multi-species methods such as bTSSFinder [2] and G4PromFinder [4]. Additionally, we included BPROM [3] in the comparative assessment, as it is the most commonly used promoter prediction program (as per Google Scholar, BPROM’s manuscript has been cited 547 times). Finally, we also compared Promotech’s performance with two recent *E. coli* -specific methods: MULTiPly [8], designed for various sigma factors, and iPro70-FMWin [14], designed for sigma 70. In total, this benchmark included Promotech’s six models and five other bacterial promoter prediction tools.

Promotech’s random forests models (RF-TETRA and RF-HOT) consistently achieved the highest AUPRC and AUROC across the four bacterial species (Tables 8 and 9). RF-TETRA achieved the highest average AUPRC and AUROC among all the methods. Among the five other bacterial promoter prediction tools, iPro70-FMWin showed the best predictive performance but still substantially lower than Promotech’s. Based on these results, we selected RF-HOT as Promotech’s predictive model for genome-wide promoter prediction. For recognizing promoters on 40-nt long genomic sequences in datasets containing up to thousands of sequences, we recommend both RF-HOT and RF-TETRA models.

**Table 8.** AUPRC per bacterial species and mean AUPRC ± standard deviation for each model. AUPRC is roughly the weighted average precision across all recall levels. A perfect classifier has an AUPRC of 1; while a random classifier has an AUPRC of 0.5 in a balanced data set. These results were obtained in balanced datasets (i.e., with a 1:1 ratio of positive to negative instances). The numbers in bold indicate the model with the highest AUPRC. For BPROM, bTSSFinder and G4PromFinder, the numbers between brackets indicate precision and recall achieved as these tools did not provide a probability associated to each instance in the dataset.

| Model | *M. smegmatis* | *L. phytofermentans* | *B. amyloliquefaciens* | *R. capsulatus* | Mean AUPRC |
| --- | --- | --- | --- | --- | --- |
| RF-HOT | **0.955** | **0.626** | 0.608 | **0.691** | 0.720±0.161 |
| RF-TETRA | 0.800 | 0.608 | **0.843** | 0.678 | **0.732±0.108** |
| GRU-0 | 0.646 | 0.486 | 0.486 | 0.588 | 0.552±0.079 |
| GRU-1 | 0.622 | 0.490 | 0.500 | 0.576 | 0.547±0.063 |
| LSTM-3 | 0.625 | 0.499 | 0.494 | 0.559 | 0.544±0.061 |
| LSTM-4 | 0.623 | 0.501 | 0.505 | 0.573 | 0.550±0.059 |
| MULTiPly | 0.649 | 0.474 | 0.653 | 0.591 | 0.592±0.083 |
| iPro70-FMWin | 0.652 | 0.582 | 0.774 | 0.594 | 0.65±0.088 |
| bTSSFinder | (0.512, 0.272) | (0.507, 0.944) | (0, 0) | (0.513, 0.250) | NA |
| G4PromFinder | (0.506, 0.938) | (0.448, 0.216) | (0.382, 0.339) | (0.510, 0.960) | NA |
| BProm | (0.781, 0.006) | (0.501, 0.560) | (0.701, 0.421) | (0.615, 0.011) | NA |

**Table 9.** AUROC per bacterial species in the validation dataset and mean AUROC ± standard deviation for each model. AUROC is roughly the likelihood that a positive instance will get a higher probability of being a promoter sequence than a negative instance. These results were obtained in datasets (not seen during training) with a 1:1 ratio of positive to negative instances. The numbers in bold indicate the model with the highest AUROC. For BPROM, bTSSFinder and G4PromFinder, the numbers between brackets indicate True Positive Rate and False Positive Rate obtained as these tools did not provide a probability associated to each instance in the dataset.

| Model | *M. smegmatis* | *L. phytofermentans* | *B. amyloliquefaciens* | *R. capsulatus* | Mean AUROC |
| --- | --- | --- | --- | --- | --- |
| RF-HOT | **0.939** | 0.591 | 0.640 | 0.660 | 0.708±0.157 |
| RF-TETRA | 0.814 | **0.608** | **0.837** | **0.674** | **0.733±0.110** |
| GRU-0 | 0.630 | 0.488 | 0.496 | 0.577 | 0.548±0.068 |
| GRU-1 | 0.601 | 0.487 | 0.502 | 0.566 | 0.539±0.054 |
| LSTM-3 | 0.622 | 0.489 | 0.481 | 0.553 | 0.536±0.066 |
| LSTM-4 | 0.592 | 0.498 | 0.506 | 0.546 | 0.536±0.043 |
| MULTiPly | 0.684 | 0.470 | 0.700 | 0.593 | 0.612±0.106 |
| iPro70-FMWin | 0.642 | 0.587 | 0.779 | 0.575 | 0.646±0.093 |
| bTSSFinder | (0.272, 0.265) | (0.944, 0.924) | (0, 0) | (0.250, 0.245) | NA |
| G4PromFinder | (0.938, 0.932) | (0.216, 0.269) | (0.339, 0.554) | (0.960, 0.953) | NA |
| BProm | (0.006, 0.002) | (0.560, 0.398) | (0.421, 0.181) | (0.011, 0.007) | NA |

Additionally, we compared the performance of RF-TETRA and RF-HOT for identifying *E. coli* promoters against that of *E. coli* -specific tools. To do this, we obtained Promotech’s predictions on a balanced data set with 2,860 experimentally validated *E. coli* promoters collected from RegulonDB [26]. This data set has been used to evaluate the performance of several *E. coli* promoter prediction tools [10, 12, 27]. The average 5-fold cross-validation MCC and accuracy reported on this data set [10, 12, 27] are in the range of [0.498, 0.763] and [0.748, 0.882], respectively. RF-HOT achieved on this data set a MCC of 0.54, accuracy of 0.77, AUPRC of 0.845 and AUROC of 0.84; while, RF-TETRA achieved a MCC of 0.47, accuracy of 0.734, AUPRC of 0.830 and AUROC of 0.808. Thus, RF-HOT is in the range of performance level observed in programs specifically developed to identify *E. coli* promoters. These results demonstrate that Promotech is indeed suitable for predicting promoters on various bacterial species.

## Conclusions

Based on our results, we recommend 1) to use *E. coli* - specific tools to predict *E. coli* promoters as they can identify *E. coli* promoters more accurately than a general bacterial promoter identification method such as Promotech; and, 2) to use Promotech to identify promoters in bacterial species other than *E. coli*, as we have shown Promotech outperforms other promoter prediction tools including iPro70-FMWin, one of the most accurate *E. coli* -specific tools [13], for identifying promoters on a variety of bacterial species (Tables 8 and 9).

In sum, Promotech is a promoter recognition tool suited for general (species independent) bacterial promoter detection that is able to perform promoter recognition on a whole bacterial genome. Promotech is available under the GNU General Public License at https://github.com/BioinformaticsLabAtMUN/PromoTech (DOI: 10.5281/zenodo.4737459).

## Methods

The goal of this study was to develop a general tool to recognize bacterial promoters. To do this, we assembled a large dataset of promoter sequences from various bacterial species, generated twelve machine learning models, and selected the best models based on AUROC and AUPRC. Our best models were compared with five existing tools (BPROM [3], bTSSFinder [2], MULTIPly [8], iPro70-FMWin [14] and G4PromFinder [4]) using a validation data set, not used for training, of four bacterial species.

### Materials

#### Collecting data

Bacterial TSS detected by next-generation sequencing (NGS) approaches, namely, dRNA-seq [15] and Cappable-seq [16], were collected from the literature (Supplementary Tables 2). We obtained promoter genomic coordinates and the corresponding sequences using BEDTools [28]. E*σ* covers DNA from roughly 55 bp upstream to 15 bp downstream of the TSS [1]. As the promoter region is not located downstream of the TSS and E*σ* covers 15bp downstream of the TSS, we excluded 15 bp from both sides of the region covered by E*σ*, and took as the promoter sequence the 40bp upstream of the TSS. The bacterial species included in this study are listed in Supplementary Table 2.

#### Generating positive and negative instance sets

A Nextflow [29] pipeline was designed to obtain the promoter (positive) sequences from the TSS coordinates by taking the genome FASTA file and the TSS coordinates as input, and obtaining the promoter coordinates as 40 bp upstream from the TSS to the TSS using BEDTools’ slop command. The BEDTools’ subtract command was used to delete duplicates, and the getfasta command was used to obtain the FASTA sequences from the promoter coordinates.

To obtain non-promoter (negative) sequences, we used BEDTools’ random command to obtain random genomic coordinates, and the getfasta command to obtain the corresponding genomic sequences. Negative sequences overlapping positive sequences were excluded from the training data. Note that some of these negative instances might in fact be actual promoters, and thus predictive performance is conservatively assessed.

The training data sets created have a 1:10 ratio of positive to negative instances (unbalanced). For the validation data set, we created a data set with a 1:1 ratio of positive to negative instances (balanced). The total number of positive and negative instances per bacterium are shown in Supplementary Tables 3 and 4 in Additional File 1.

### Machine learning models

We used two machine learning methods: Recurrent Neural Networks (RNNs) [19] and Random Forest (RF) [17, 18]. Both methods have been successfully used before to classify genomic sequences. Random Forest is a popular machine learning method for its ability to identify feature importance, handles many data types (continuous, categorical and binary). It is well-suited for high-dimensional data, and avoids overfitting by its voting-scheme among the ensemble of trees within it [30]. RNNs are also well-suited for genomic sequence analysis due to their ability to handle variable-length inputs, detecting sequential patterns, and retaining information through time.

#### Model selection

Due to the lengthy training time of the RNNs, we were unable to run CV for the RNN models. Thus, to select the best model, we split our training data in 75% for training and 25% for testing. Models were trained with 75% of the data, and then compared to each other on their performance in the 25% left-out data. After selecting the best models, these models were retrained using all of the training data, and the resulting models used for whole-genome promoter prediction and comparative assessment with the other tools.

#### Random forest

Two RF models were generated, the first was trained using hot-encoded features; this meant that the nucleotides (A, G, C, T) were transformed into binary vector representations [1000], [0100], [0010], and [0001], respectively. This model is henceforth referred to as RF-HOT. The second model was trained using tetra-nucleotide frequencies calculated using the scikit-bio library [31] and denoted as RF-TETRA. The models were created using the Sklearn’s RandomForest-Classifier [32] combined with a 10-fold grid search CV to handle the hyper-parameter optimization. The hyper-parameter search space was max features (m): [None, “sqrt”, “log2”] and n estimators (n): [1000, 2000, 3000]; both models were trained using an unbalanced data set with a 1:10 ratio of positive to negative instances. The best hyper-parameters found by grid-search CV for both RF models were m=“log2” and n=2,000 with class weights values of *{*0 : 0.53, 1 : 10.28*}*.

#### Recurrent neural networks

Two types of RNNs were trained, Long Short Term Memory Unit (LSTM) [20] and Gated Recurrent Unit (GRU) [21], using word embeddings representation of the tetra-nucleotide sequences calculated using the Keras’ Tokenizer class [33]. The models were designated as GRU-X or LSTM-X where X indicates the number of hidden layers. All models were manually tuned with an architecture consisting of one embedding layer, one GRU or LSTM layer, zero to four dense layers with dropout to reduce overfitting, one binary output, Adam optimizer function and Binary Crossentropy loss function (Table 5 in Additional File 1).

#### Computer infrastructure

All RF and RNN models were trained on the Compute Canada’s Beluga Cluster [34] configured with four NVidia V100SXM2 16GB GPUs, eight Intel Gold 6148 Skylake @ 2.4 GHz CPUs, and managed using SLURM commands.

### Model assessment

Three assessments were performed to evaluate models’ performance. The first consisted in scanning each bacterial genome using a 40-nt sliding window. In total, the number of generated sequences ranged from 4 to 7 million depending on the genome size. Models were given as input each of the sliding window sequences. Models then outputted per 40-nt sequence the probability of having a promoter within that sequence. To be counted as a true positive, the predicted promoter sequence had to have at least 10% sequence overlap with an actual promoter sequence. All other predicted promoters were considered false positives. To determine whether a predicted promoter overlapped with an actual promoter, we used the BEDTools intersect command with the parameters -s and -f 0.1.

In the second assessment, we also scanned each bacterial genome using a 40-nt sliding window. However, in this assessment we considered a predicted promoter a true positive if it was within 100 nt of an actual promoter. In this setting, we use BEDtools’ closest command to find the five predicted promoters closest to an actual promoter. Then, those closest predicted promoters less than 100 nt away, upstream or downstream, from an actual promoter were counted as true positives. All other predicted promoters were considered false positives.

In the third assessment we used the validation balanced data set obtained as described above. In this assessment we included BPROM, bTSSfinder, G4Promfinder, iPro70-FMWin, and MULTiPLy. MUL-TiPLy and G4PromFinder accept 40 nt-long sequences, but BPROM and bTSSFinder require sequences 250-nt long and iPro70-FMWin requires 81-nt long sequences. We used BEDTools’ slopBed [38] command to extend the sequences in the dataset from 40 to the required length. As G4Promfinder is written in Python, we integrated into our own pipeline and fed the sequences directly to G4Promfinder. To run, BPROM and bTSSfinder we wrote each sequence to a file and then ran the programs through a shell script called from our own pipeline. MULTiPLy was tested separately, as it was developed in Matlab, so a short script was written to feed it each bacterium’s data set. We ran BPROM and MULTiPLy with their default values. bTSSFinder was run with the parameters -c 1 -t e and -h 2 indicating to search for the highest ranking promoter regardless of promoter class, use *E. coli* mode, and search on both strands. All other bTSSFinder parameters were left at their default values. G4Promfinder only requires as input sequences in FASTA format. iPro70-FMWin’s results were obtained from its web site inputting the sequences in FASTA format.

Performance metrics used were: 1) the area under the precision-recall curve (AUPRC), where precision is the number of true positives divided by the total number of predicted positives and recall (also called sensitivity or true positive rate) is the number of true positives divided by the total number of actual positives; and 2) the area under the ROC curve (AUROC), where true positive rate is the same as recall and false positive rate is the number of false positives divided by the total number of predicted positives.

## Declarations

Ethics approval and consent to participate Not applicable.

Consent for publication Not applicable.

Availability of data and materials

All data, scripts and pipelines used are available in https://github.com/BioinformaticsLabAtMUN/PromoTech and https://doi.org/10.5281/zenodo.4737459.

## Author’s contributions

L.P.-C. conceived Promotech. R.C. implemented Promotech and performed all the experiments under the supervision of L.P.-C. All authors discussed the results and contributed to the final manuscript.

## Acknowledgements

This research was enabled in part by support provided by ACENET (www.ace-net.ca/) and Compute Canada (www.computecanada.ca).

## Additional Files

Additional File 1 — The Additional File 1 is a PDF file that includes the following tables:

1. **Supplementary Table 1** — A summary of promoter prediction approaches in the last twelve years.
2. **Supplementary Table 2** — A summary of all the bacteria used during their study and their properties.
3. **Supplementary Table 3** — The number of TSS per each bacterium in the training data set.
4. **Supplementary Table 4** — The number of TSS per each bacterium in the validation data set.
5. **Supplementary Table 5** — RNN hyper-parameters.
6. **Supplementary Figs. 1 - 4** — PRC and ROC per model and per bacterium when requiring 10% sequence overlap between predicted promoters and actual promoters to count them as true positives.

