## Supplementary Tables and Figures for "Promotech: A general tool for bacterial promoter recognition"

### Promotech's additional file

Ruben Chevez-Guardado and Lourdes Peña Castillo
Memorial University of Newfoundland

April 30, 2021

#### Contents

**1 Computational approaches for bacterial promoter recognition 2**

**2 Data sets used in this study 4**

**3 Whole-genome promoter search requiring 10% sequence overlap between predicted promoters and actual promoters 9**

**4 RNN's hyper-parameters 11**

### 1 Computational approaches for bacterial promoter recognition

Table 1: A number of promoter prediction approaches published within the last twelve years. Columns are methods' name and publication year, brief description of approach used to detect promoters, and target organism(s).

| Name | Approach | Target Organism(s) |
|---|---|---|
| Selector [1] (2021) | Stacked-ensemble approach | *E. coli* |
| iPromoter-BnCNN [2] (2020) | CNN-based classifier | *E. coli* |
| MULTiPLy [3] (2019) | SVM | *E. coli* |
| iProEP [4] (2019) | Support vector machine (SVM) | *H. sapiens*<br>*D. melanogaster*<br>*C. elegans*<br>*B. subtilis*<br>*E. coli* |
| iPro70-FMWin [5] (2018) | Sequence-based features extracted for "multiple windows" and logistic regression | *E. coli* |
| IBPP [6] (2018) | SVM | *E. coli* |
| iPromoter-2L [7] (2018) | Multi-window-based pseudo K-tuple nucleotide composition | *E. coli* |
| 70ProPred [8] (2018) | SVM | *E. coli* |
| G4PromFinder [9] (2018) | Detection of AT-rich element and G-quadruplex motifs | *S. coelicolor A3(2)*<br>*P. aeruginosa PA14* |

| | | |
|---|---|---|
| bTSSfinder [10] (2017) | Position weight matrices (PWM) and neural networks | *E. coli K12*<br>*Nostoc sp. PCC 7120*<br>*Synechocystis sp. PCC 6803*<br>*S. elongatus PCC 6301* |
| CNNProm [11] (2017) | Convolutional Neural Networks (CNN) | *E. coli*<br>*B. subtilis*<br>*Homo sapiens*<br>*Mus musculus*<br>*Arabidopsis* |
| vw Z-curve [12] (2012) | Variable-window Z-curve for frequencies of $k$-nucleotides and partial least squares classifier | *E. coli*<br>*B. subtilis* |
| PePPER [13] (2012) | MEME motif search for the Pribnow box DNA pattern | *L. lactis*<br>*E. coli*<br>*B. subtilis* |
| BacPP [14] (2011) | Neural networks | *E. coli* |
| PromPredict [15] (2010) | Thresholds on sequences' average free energy | *E. coli*<br>*B. subtilis*<br>*M. tuberculosis* |
| BPROM [16] (2010) | Linear Discriminant Analysis (LDA) | *E. coli* |

### 2 Data sets used in this study

Table 2: Summary of data sets used. In the last column a T or V indicates whether the bacterium is reserved for training or validation, respectively. Additional information is included such as the number of TSS per bacterium, the genome's length, the Next Generation Sequencing technology used to obtain the TSSs, and the literature sources' PubMed ID ( if PubMed ID is missing then, that at the time of this publication, the source manuscript was still in preparation.

| BACTERIUM | GENOME ACCESSION | PUBMED-ID | NGS TECHNOLOGY | #TSS | GENOME LENGTH | T or V |
|---|---|---|---|---|---|---|
| *Escherichia coli* str. K-12 substr. MG1655 | NC_000913.3 | 27748404 [17] | dRNA-seq | 278 | 4,641,652 | T |
| *Escherichia coli* str. K-12 substr. MG1655 | NC_000913.2 | 25266388 [18] | dRNA-seq | 2,672 | 4,639,675 | T |
| *Helicobacter pylori* 26695 | NC_000915.1 | 20164839 [19] | dRNA-seq | 1,907 | 1,667,867 | T |
| *Helicobacter pylori* 26695 | NC_000915.1 | 30169674 [20] | dRNA-seq | 449 | 1,667,867 | T |
| *Campylobacter jejuni subsp. jejuni* 81116 | NC_009839.1 | 30169674 [20] | dRNA-seq | 269 | 1,628,115 | T |
| *Campylobacter jejuni subsp. jejuni* NCTC 11168 | NC_002163.1 | 23696746 [21] | dRNA-seq | 1,905 | 1,641,481 | T |
| *Campylobacter jejuni* RM1221 | NC_003912.7 | 23696746 [21] | dRNA-seq | 2,167 | 1,777,831 | T |
| *Campylobacter jejuni subsp. jejuni* 81116 | NC_009839.1 | 23696746 [21] | dRNA-seq | 1,944 | 1,628,115 | T |

| | | | | | | |
|---|---|---|---|---|---|---|
| *Campylobacter jejuni subsp. jejuni* 81-176 | NC_008787.1 | 23696746 [21] | dRNA-seq | 2,003 | 1,616,554 | T |
| *Streptococcus pyogenes* strain S119 | LR031521.1 | 30902048 [22] | dRNA-seq | 892 | 1,877,450 | T |
| *Salmonella enterica subsp. enterica serovar Typhimurium* SL1344 | NC_016810.1 | 22538806 [23] | dRNA-seq | 1,873 | 4,878,012 | T |
| *Chlamydia pneumoniae* CWL029 | NC_000922.1 | 21989159 [24] | dRNA-seq | 530 | 1,230,230 | T |
| *Shewanella oneidensis* MR-1 | NC_004347.2 | 24987095 [25] | dRNA-seq | 4,729 | 4,969,811 | T |
| *Leptospira interrogans serovar Manilae* isolate L495 | NZ_LT962963.1 | 28154810 [26] | dRNA-seq | 2,865 | 4,614,703 | T |
| *Streptomyces coelicolor* A3(2) | NC_003888.3 | 27251447 [27] | dRNA-seq | 3,570 | 8,667,507 | T |
| *Mycolicibacterium smegmatis* str. MC2 155 | NC_008596.1 | 30984135 [28] | dRNA-seq | 4,054 | 6,988,209 | V |
| *Lachnoclostridium phytofermentans* ISDg | NC_010001.1 | 27982035 [29] | Cappable-seq | 1,187 | 4,847,594 | V |
| *Rhodobacter capsulatus* SB 1003 | NC_014034.1 | - [30] | dRNA-seq | 5,374 | 3,738,958 | V |

| | | | | | | |
|---|---|---|---|---|---|---|
| *Bacillus amyloliquefaciens* XH7 | CP002927.1 | 26133043 [31] | dRNA-seq | 1,064 | 3,939,203 | V |

Table 3: Training data sets. Data sets are given in the same order as in Table 2

| BACTERIA ID | PROMOTER SEQUENCES | NON-PROMOTER SEQUENCES |
|---|---|---|
| ECOLI | 248 | 2,773 |
| ECOLI_2 | 2,636 | 26,147 |
| HPYLORI | 1,877 | 18,273 |
| HPYLORI_2 | 448 | 4,448 |
| CJEJUNI | 269 | 2,674 |
| CJEJUNI_2 | 1,881 | 18,298 |
| CJEJUNI_3 | 2,140 | 20,736 |
| CJEJUNI_4 | 1,919 | 18,619 |
| CJEJUNI_5 | 1,973 | 19,173 |
| SPYOGENE | 891 | 8,754 |
| STYPHIRMURIUM | 1,869 | 18,464 |
| CPNEUMONIAE | 530 | 5,221 |
| SONEIDENSIS | 4,728 | 46,511 |
| LINTERROGANS | 2,791 | 27,979 |
| SCOELICOLOR | 3,566 | 35,170 |
| **SUBTOTAL** | 27,766 | 273,240 |
| **TOTAL** | 301,006 | |

Table 4: Balanced test data sets. These data sets were not used during training. They were used to assess Promotech's performance on independent data and compare its performance with that of existing tools.

| BACTERIA ID | PROMOTER SEQUENCES | NON-PROMOTER SEQUENCES |
|---|---|---|
| MYCOBACTER | 4,054 | 3,978 |
| CLOSTRIDIUM | 1,187 | 1,177 |
| RHODOBACTER | 5,374 | 5,207 |
| BACILLUS | 1,064 | 1,055 |
| **SUBTOTAL** | 11,615 | 11,417 |
| **TOTAL** | 23,032 | |

### 3 Whole-genome promoter search requiring 10% sequence overlap between predicted promoters and actual promoters

Figure 1: A comparison between the AUPRC and AUROC requiring 10% sequence overlap between predicted promoters and actual promoters in the *Mycolicibacterium smegmatis* str. MC2 155 bacterium. Column **(a)** shows the PR curves per model and **(b)** shows the ROC curves.

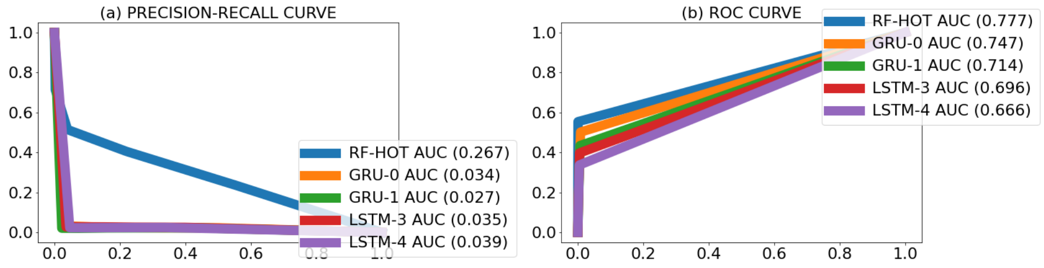


Figure 2: A comparison between the AUPRC and AUROC requiring 10% sequence overlap between predicted promoters and actual promoters in the *Lachnoclostridium phytofermentans* ISDg bacterium. Column **(a)** shows the PR curves per model and **(b)** shows the ROC curves.

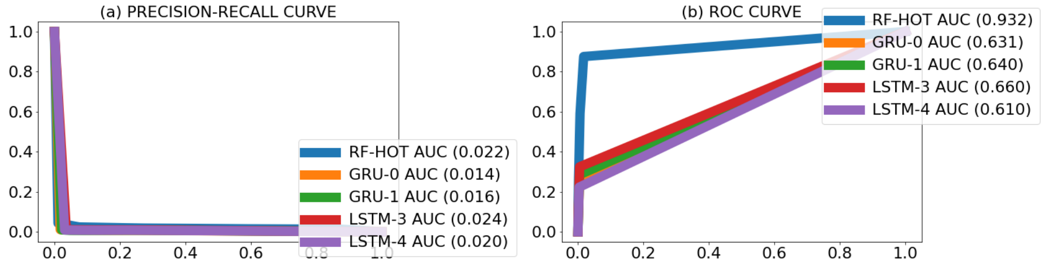

Figure 3: A comparison between the AUPRC and AUROC requiring 10% sequence overlap between predicted promoters and actual promoters in the *Rhodobacter capsulatus* SB 1003 bacterium. Column **(a)** shows the PR curves per model and **(b)** shows the ROC curves.

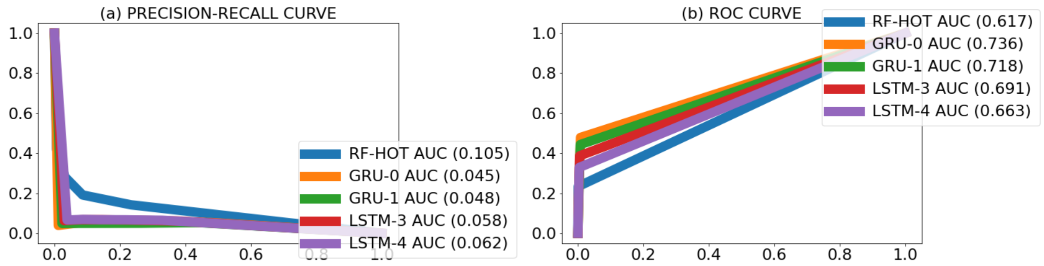


Figure 4: A comparison between the AUPRC and AUROC requiring 10% sequence overlap between predicted promoters and actual promoters in the *Bacillus amyloliquefaciens* XH7 bacterium. Column **(a)** shows the PR curves per model and **(b)** shows the ROC curves.

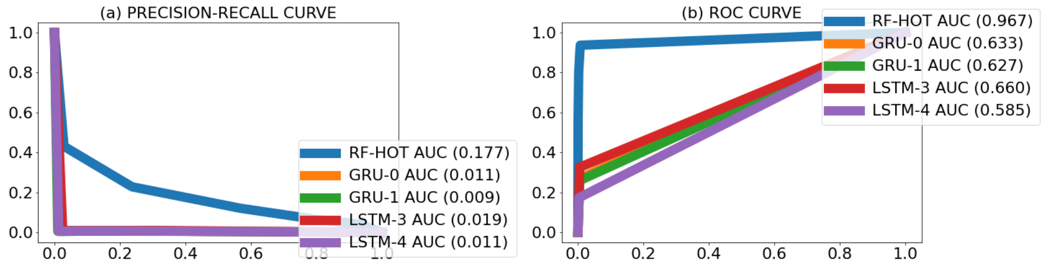

### 4 RNN's hyper-parameters

Table 5: Hyper-parameter setting used in the RNN's architecture and training.

| **Hyper-parameter** | **Value** |
|---|---|
| Number of embedding units | 50 |
| Number of recurrent units | 100 |
| Type of recurrent units | LSTM or GRU |
| Number of fully-connected units | 100 |
| Number of hidden dense layers | 0, 1, 2, 3, or 4 |
| Dropout rate | 0.2 |
| Epochs | 50 |
| Batch size | 10 |
| Activation function | Sigmoid |
| Optimizer | Adam |
| Loss function | Binary cross-entropy |

[28] M Carla Martini, Ying Zhou, Huaming Sun, and Scarlet Sara Shell. Defining the transcriptional and post-transcriptional landscapes of *My-*

*cobacterium smegmatis* in aerobic growth and hypoxia. *Frontiers in microbiology*, 10:591, 2019.

[29] Magali Boutard, Laurence Ettwiller, Tristan Cerisy, Adriana Alberti, Karine Labadie, Marcel Salanoubat, Ira Schildkraut, and Andrew C Tolonen. Global repositioning of transcription start sites in a plant-fermenting bacterium. *Nature communications*, 7(1):1–9, 2016.

[30] Marc Grüll. *Transcriptomic studies of the bacterium Rhodobacter capsulatus.* PhD thesis, Memorial University of Newfoundland, 2019.

[31] Yuling Liao, Lianggang Huang, Bin Wang, Feng Zhou, and Li Pan. The global transcriptional landscape of *Bacillus amyloliquefaciens XH7* and high-throughput screening of strong promoters based on RNA-seq data. *Gene*, 571(2):252–262, 2015.
